# Proteome integral solubility alteration assay for monitoring antibody-antigen cell interactions

**DOI:** 10.1101/2025.09.04.673944

**Authors:** Weiqi Lu, Susanna L. Lundstrom, Yaolin Hou, Sourav Mukherjee, Jesper Z. Haeggström, Massimiliano Gaetani, Roman A. Zubarev

**Author notes:** These authors contributed equally to this work (Weiqi Lu, Susanna L. Lundstrom).

## Abstract

The Proteome Integral Solubility Alteration (PISA) assay is commonly used to identify and characterize interactions between small molecules and the cellular proteome. In this study, a modified PISA approach identified tumor necrosis factor alpha (TNFα) as the antigen of the monoclonal antibody infliximab in activated monocytes - uniquely among ∼8,000 cellular proteins. These results highlight the potential of PISA-based analysis not only for monitoring antibody-antigen interactions in therapeutic development, but also for identifying new antigens and characterizing the initial effector functions of antibodies within complex proteome environments.

**Teaser:** Modified PISA pinpointed TNF-α as Infliximab’s antigen among 8000 proteins, highlighting broad potential for therapeutic antigen studies.

## Introduction

Monitoring drug–target interactions in vitro and in vivo is critical in drug development. The Proteome Integral Solubility Alteration (PISA) assay, which detects changes in protein solubility within the proteome matrix upon drug binding, has become a widely used method for monitoring protein–drug interactions, particularly in drug target deconvolution studies (1-5). While most approved pharmaceuticals are small molecules, the rapid rise of antibody therapeutics (6) and the emerging field of intracellular functional antibody delivery (7, 8) emphasize the need for tools capable of monitoring antibody–antigen interactions. However, adapting PISA to monoclonal antibodies (mAbs) presents unique technical challenges, including issues related to their large size and high unfolding temperatures. In this study, we optimized a PISA workflow to monitor antibody–antigen interactions and successfully identified the cytokine tumor necrosis factor alpha (TNFα) as the antigen of the therapeutic mAb infliximab (INX) in human monocyte lysates (ML) and intact monocytes.

TNFα is a cytokine that modulates innate immune inflammatory reactions as well as cellular activation leading to proliferation, programmed cell death and necrosis (9-12). In macrophages and monocytes, the production of TNFα can be triggered by lipopolysaccharides (LPS) (13). INX is a chimeric mAb that is binding both monomeric and trimeric forms of TNF-α (14, 15). Besides its own affinity to soluble (s)TNF-α, INX binds its precursor, membrane (m)TNF-α, which also participates in cell signaling (13). Given the well-characterized properties and accessibility of both the antibody and its antigen, they were well suited for the PISA pilot experiments. To distinguish direct antibody–antigen interactions from antibody cross-reactivity and effector-related activities, the experiments were further conducted in parallel using both intact antibody and the Fab fragment.

This proof-of-concept pilot demonstrates that PISA can be extended beyond small-molecule interactions to study complex antibody–antigen binding in proteome-wide environments. An overview of the optimized workflow is presented in **Figure 1**.

**Figure 1.**
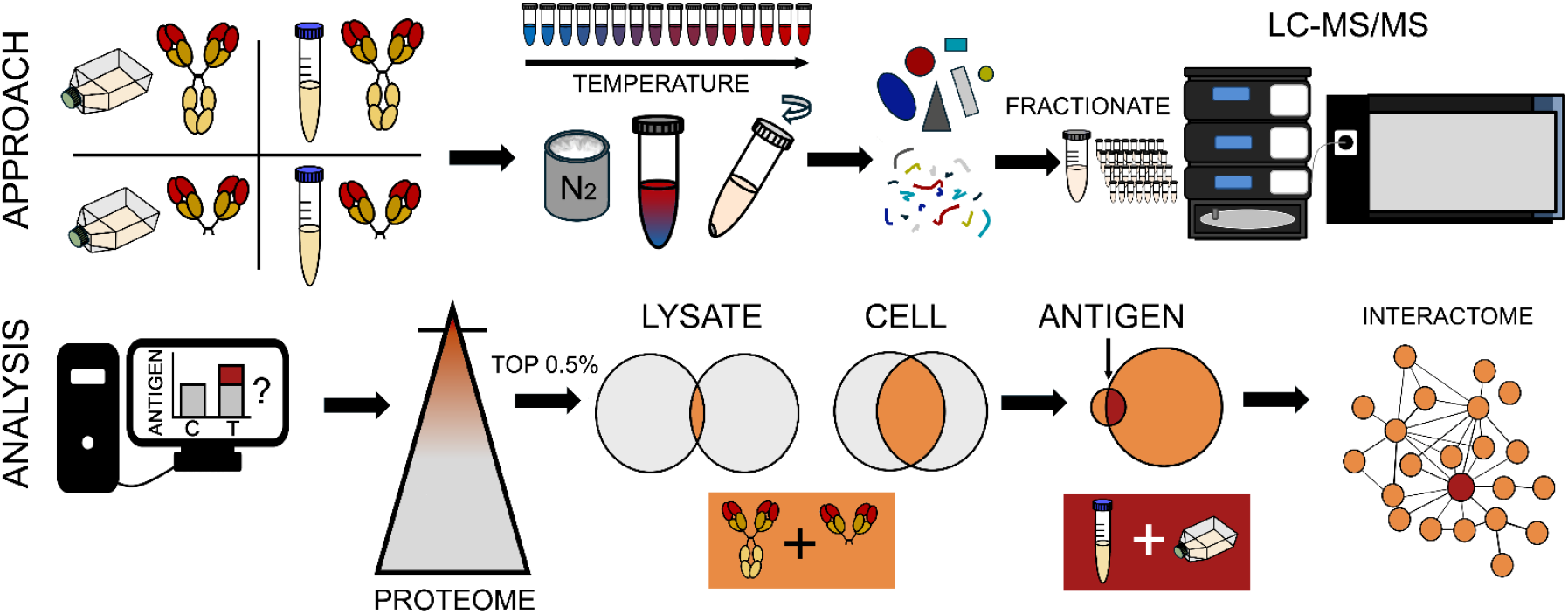
Overview of the optimized PISA workflow for antibodies. Cell lysates and/or live cells are treated with either the intact antibody or the Fab fragment, alongside appropriate controls. Samples are aliquoted, subjected to temperature treatment, snap-frozen in liquid nitrogen, pooled, lysed (for intact cells), and ultracentrifuged. The resulting supernatants are analyzed by proteomics using TMT labeling. For data analysis, control and antibody-treated samples are compared to determine which proteins show the greatest solubility shifts between conditions. The top 0.5% of proteins with the largest and most significant shifts across all four experimental setups (lysate - antibody-treated, lysate - Fab-treated, intact cell - antibody-treated, intact cell - Fab-treated) are selected. These proteins are then compared to identify both the antigen and the broader initial antibody interactome.

## Results

### Proof-of-principle experiment in control lysates

We first performed PISA on control human monocyte lysate spiked with tumor necrosis factor alpha (TNFα). The experiment was performed using either intact INX or Fab-INX, respectively. We used a modified PISA scoring system (4, 16, 17) to identify the most probable antigen candidate in the experiments. Thus, the score for each protein was calculated as the absolute value of the logarithm of the PISA signal ratio with and without either INX or its Fab region (Fab-INX) multiplied by the statistical significance -log10(p). TNFα was identified by twelve unique peptides and ranked, on average, as approximately the 1800th most abundant protein in the dataset. Despite its moderate abundance, it emerged as the top candidate for both intact INX and Fab-INX among 4597 and 6365 quantified proteins, respectively (**Figure 2A–B**). Highlighting the complex nature of protein solubility, the PISA signal of TNFα exhibited a nonlinear shift upon the addition of mAb to cell lysates with increasing total protein concentration—changing sign twice in the process. This behavior remained largely the same when the cell lysate was replaced by human serum albumin (**Figure 2C**). Such behavior complicated the statistical analysis, which was ameliorated by the use of Fisher’s formula (18) for calculating the p-value of each protein from its three most statistically significantly changing peptides.

**Figure 2.**
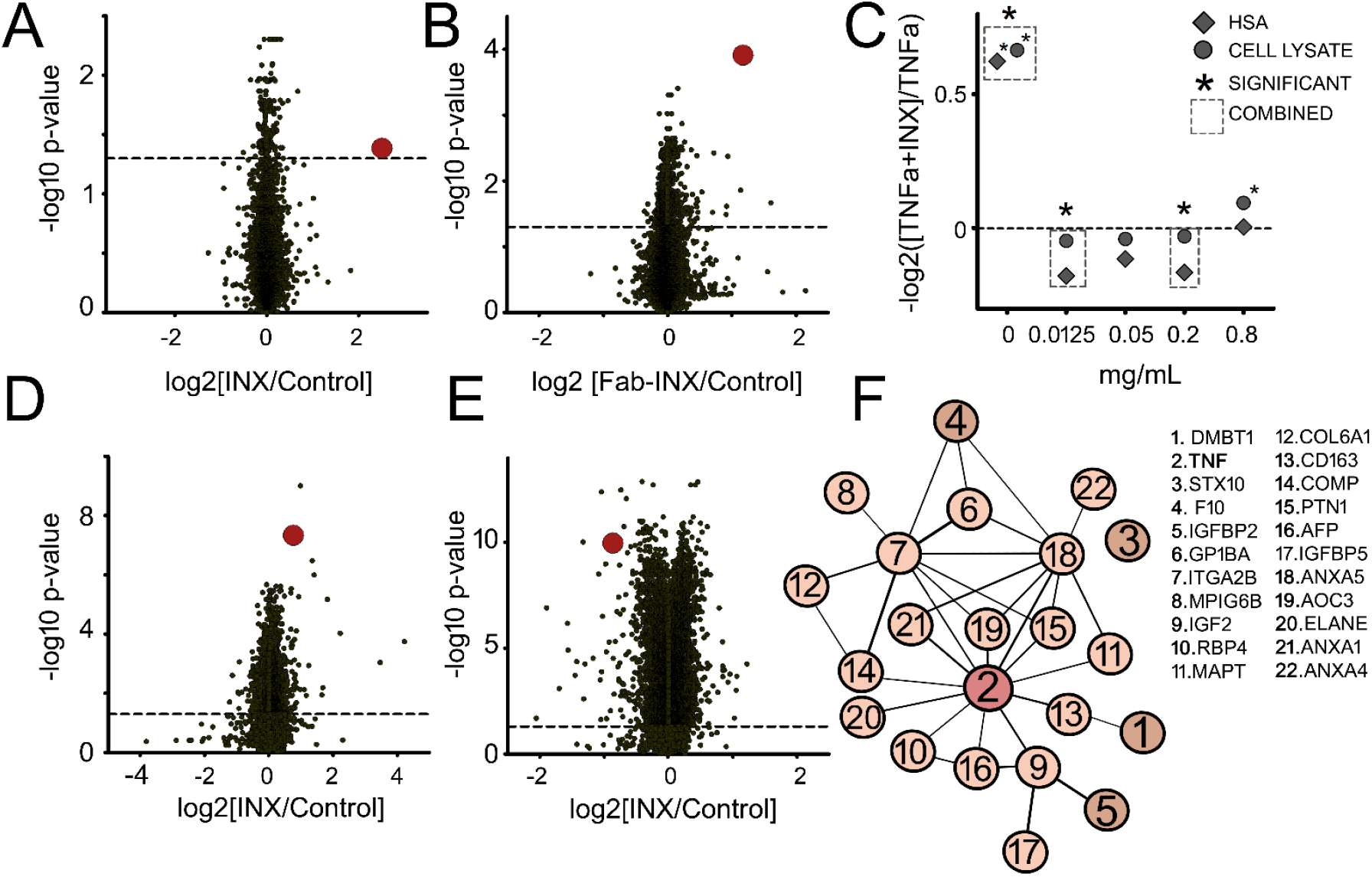
PISA-AAI, summary of results. (**A**) Volcano plot from the experiment using control human monocyte lysate spiked with TNFα, comparing intact antibody (INX) to control. TNFα is shown as a larger red circle. (**B**) Volcano plot from the experiment using control human monocyte lysate spiked with TNFα, comparing the Fab fragment (Fab-INX) to control. TNFα is shown as a larger red circle. (**C**) Relative change in TNFα levels between samples containing TNFα + INX versus TNFα alone under varying background protein concentrations (0–0.8 mg/mL) of either HSA or cell lysate. (**D**) Volcano plot from the experiment using LPS-treated monocyte lysate, comparing solubility shifts between INX and control. TNFα is shown as a larger red circle. (**E**) Volcano plot from the experiment using LPS-treated intact monocytes subjected to PISA, comparing solubility shifts between INX and control. TNFα is shown as a larger red circle. (**F**) STRING pathway analysis of the 22 overlapping proteins identified among the top 0.5% in both Fab and intact antibody experiments in LPS-treated monocytes.

### Endogenous TNFα targeting in LPS-stimulated monocytes

We next analyzed lysates from LPS-stimulated monocytes derived from human blood and quantified endogenous TNFα using ten unique peptides (**Supplemental Table 1**). TNFα ranked as the 1445th most abundant protein among 7794 quantified proteins. Upon addition of either INX or Fab-INX to the lysate, TNFα solubility increased significantly, ranking #8 with INX (**Figure 2D**) and #2 with Fab-INX. Among the top 0.5% of candidate targets (39 proteins) in both datasets, only TNFα and telomerase-associated protein 1 (TEP1)—ranked 31st with INX and 5th with Fab-INX— were common (**Supplemental Table 2**). TNFα had a substantially better average rank (5 vs. 18), making it the strongest overall antigen candidate. Noteworthy, many of the top proteins exclusively present in the INX dataset (e.g., IgM, IgA, FCRL3, ALB, TF, APOA1, APOA2, A2M, ORM1, FCRL3, IGHM and IGHA1, etc.) are known to interact with the IgG Fc-region that is absent in Fab-INX. This finding opens way for filtering out unspecific interactions of proteins with antibodies.

### Endogenous antigen targeting in intact monocytes

Finally, PISA was performed in LPS treated, intact human monocytes. TNFα was quantified with 3 unique peptides (**Supplemental Table 1**), ranking as 2395-th most abundant among the 8134 quantified proteins. TNFα solubility significantly decreased when antibody or its Fab region were added. TNFα scored as #8 for INX (**Figure 2E**) and #13 for Fab-INX. The overlap of the top 0.5% highest ranking candidate targets (41 proteins) gave many more proteins (22) than in the cell lysate experiment, and the TNFα did not emerge as the topmost-ranking candidate in the combined list (**Supplemental Table 3**). However, upon incorporating prior insights from the lysate experiments, specifically that solubility shifts are more pronounced with full antibodies (150 kDa) than with Fab fragments (50 kDa), and that both shifts occur in the same direction, the list narrowed to just five candidate proteins. Among these, TNFα ranked second after DMBT1. However, because DMBT1 is a scavenger receptor cysteine-rich (SRCR) domain–containing protein known to bind non-specifically to immunoglobulins (19, 20), it can be considered a false positive. Notably, TNFα was the only protein common between the two (INX / Fab-INX)-overlapping proteins in lysate and 22 such overlapping proteins in intact cells. Taken together, these findings uniquely identified TNFα as the antigen targeted by INX in intact human monocytes.

The significantly larger number of overlapping proteins between INX and Fab-INX conditions in intact cells, compared to lysates, is likely attributable to protein-protein interactions. Indeed, STRING pathway analysis revealed that these proteins are inter-connected and engaged in an extended network encompassing almost all (21) of these proteins (**Figure 2F**). Not surprisingly, TNF emerged as the most prominent node with twelve connecting proteins, while the two next-best connected molecules, ITGA2B and ANXA5, had nine connections each. Therefore, pathway analysis confirmed that TNF is the most important candidate. Note that upstream pathway analysis of most affected proteins can correctly identify the drug target even when the target protein itself is absent in the input list (21). Therefore, antibody-antigen interactions can be monitored within complex cellular environments through the behavior of not only the antigen itself, but also of the multitude of proteins interacting with it.

## Discussion

In this study, we demonstrate that the PISA assay can be adapted to investigate antibody– antigen interactions (PISA-AAI) within complex proteomic environments. Using infliximab (INX) as a model antibody, we show that PISA-AAI consistently identified tumor necrosis factor alpha (TNFα) as the antigen, both in cell lysates and in intact monocytes, despite its only moderate abundance among the thousands of proteins detected. Importantly, the approach also captured network-level effects by revealing a broader set of proteins that shift in solubility upon antibody treatment, providing insight into the interactome surrounding the antibody–antigen complex. Together, these findings establish PISA-AAI as a functional and quantitative tool for monitoring antibody interactions directly in physiologically relevant systems.

To date, all antibody-based therapeutics target extracellular soluble proteins or membrane surface proteins. Despite the extraordinary potential of the antibody therapeutics, it has been limited by the inability of the antibody to pass through the cell membrane. In vitro, intracellular delivery of antibodies and other proteins can be achieved through electroporation or by exploiting endocytosis pathways using molecular binders such as small molecules, protein toxins, or peptides (7, 8). The emerging field of intracellular functional antibody delivery as a targeted therapeutic approach (22-24) holds immense promise for the development of novel, highly specific, and effective treatment strategies. Thus, it is evidently important to have appropriate assays and evaluation criteria to study mAb-target interactions in complex proteome environment.

Our results have several implications for antibody-based therapeutic development. Conventional techniques such as Western blotting or fluorescence microscopy are limited in their ability to assess antibody–antigen interactions in a proteome-wide and quantitative manner. By contrast, PISA-AAI detects not only direct solubility shifts of the antigen itself but also the wider proteomic context, thereby providing complementary mechanistic information. The approach will be particularly valuable in situations where antibody delivery to intracellular targets is explored, an emerging therapeutic frontier with strong potential but considerable challenges. Furthermore, the distinct patterns observed for intact versus Fab antibodies (reflecting Fc-mediated interactions and size-dependent solubility effects), underscore the ability of PISA-AAI to differentiate specific antigen recognition from non-specific background interactions, a key step toward robust target validation.

Limitations should be acknowledged. First, the solubility behavior of antibodies is strongly dependent on the protein background: non-linear shifts and even changes in the direction of solubility effects complicate statistical interpretation. Second, Fc-mediated binding generates additional candidates that must be distinguished from true antigen interactions. Third, TNFα was only moderately abundant, ranking in the mid-thousands among detected proteins, emphasizing the need for sensitive detection and careful scoring approaches. Finally, although our analysis highlights the utility of combining intact antibody and Fab fragment experiments, this parallel design increases experimental complexity and may not always be feasible.

Future work should aim to refine statistical frameworks tailored for antibody-induced solubility changes, expand validation to a broader panel of antibodies and antigens, and integrate PISA-AAI with complementary workflows such as PISA-REX for redox (25) and expression profiling. Defining robust criteria for filtering out Fc-mediated or sticky protein interactions will further enhance specificity. Such developments will help establish PISA-AAI as a routine tool for evaluating antibody behavior across different biological contexts.

Looking ahead, PISA-AAI offers a powerful means to move beyond antigen identification alone, toward a more holistic understanding of antibody effects in the cellular proteome. By capturing both direct binding events and associated network changes, PISA-AAI has the potential to become an indispensable component of preclinical antibody evaluation, enabling more confident predictions of therapeutic efficacy and mechanisms of action.

## Material and Methods

### Experimental design

Targeting the antibody antigen interaction by PISA was performed on samples with increased complexity, starting with cell lysate with the antigen added, and then subsequently applying the method on endogenously produced antigen in cell lysate and on intact cells. Experiments were performed in triplicates or quadruplicates and the potential significant shift in the antigen and antibody were then tested and compared to the rest of the proteome.

### Antibody

INX was obtained from Merck, European Pharmacopoeia Reference Standard, (Foreign Trade Commodity Code: 30021500). INX-Fab was prepared via Pierce™ F(ab’)2 Preparation Kit (Thermo Scientific) using instructions provided by the manufacturer.

### Monocytes

Human mononuclear cells (monocytes) were isolated from buffy coats obtained from adult blood donors (Karolinska University Hospital, Stockholm, Sweden) by density-gradient centrifugation using Lymphoprep as described previously (26). Monocytes (1×106 cells/mL) were then treated for 24 h with or without 100 ng/mL LPS at 37°C in 5% CO2 in RPMI-1640 medium supplemented with 10% heat-inactivated fetal bovine serum, 2 mM glutamine, 100 U/mL penicillin, and 100 μg/mL streptomycin, to obtain control and LPS-treated monocytes, respectively. At the end of the incubation period, cell culture supernatants were removed by centrifugation (1000 g). Cells were resuspended in PBS.

To obtain cell lysate, cells were lysed by performing five freeze-thaw cycles (liquid nitrogen 37°C) followed by incubation for 30 min at 4°C in PBS with 0.1% NP40, and probe sonication on ice for 2 min (3 s pulse, 30% amplitude). Supernatants were collected following centrifugation at 10,000 g for 10 min at 4°C. The control monocytes lysates (CML) and LPS-treated monocytes lysates (LML) were transferred to protein low-bind Eppendorf tubes which were stored at -80°C until further processing. To CML (0.8 mg/mL), TNFα (0.08 µM) was added and samples were incubated for 40 min at 20°C, either alone or with 0.31 µM INX, DG-INX or Fab-INX. LML samples (0.8 mg/mL) were incubated for 40 min at 20°C either alone or with 0.31 µM INX, DG-INX or Fab-INX. In experiments using intact cells, after LPS treatment, either PBS, INX or Fab-INX were added to final concentration of 0.1 mg/mL and incubated for 1 h.

### PISA analysis

Thermal treatment was performed as previously described (1, 16). Each sample was distributed into 16 aliquots that were heated at 16 temperature points from 50 to 78°C for 3 min, and then left to rest at 25°C for 6 min. Following treatment, samples were snap frozen in liquid nitrogen and thawed at RT before further treatment. The temperature point aliquots for each sample were pooled together. For the intact cells, NP-40 protein detergent (Thermo Fisher Scientific) was added to a final concentration of 0.4%, and the sample was shaken at 300 rpm for 1 h at 4°C. Five freeze-thaw cycles were then performed to obtain the cell lysate. All samples were ultracentrifuged at 126,600 g for 30 min at 4°C using an Optima XPN-80 (Beckman Coulter, USA). The supernatants were collected. For the protein gradient experiments, aliquots of 300 µL (samples: 0, 0.125 and 0.05 mg/mL), 200 µL (samples: 0.2 mg/mL) and 100 µL (samples: 0.8 mg/mL) of the supernatant was taken out and 20 µL of an internal standard (0.4 mg/mL myoglobin and 0.6 mg/mL trypsinogen) added. The samples were then prepared for proteomics analysis as described below. For the experiments on cell lysates and intact cells, the protein concentration of the supernatant following ultracentrifugation was measured using Micro BCA Protein Assay Kit, and an equal amount of protein (50 µg per sample) was used for all proteomics analyses.

Protein samples were reduced, alkylated and digested as previously described (1, 16). Briefly, samples were reduced in 8 mM dithiothreitol (55°C, 45 min) and alkylated in 25 mM iodoacetamide (25°C, 30 min in darkness). Samples were then precipitated with acetone at -20°C overnight. Then the pellet was resuspended in 8 M urea in 20 mM EPPS buffer (pH 8.2). Samples were digested with LysC (4 M urea, 20 mM EPPS, pH 8.2, 23°C, 6 h) and then with trypsin (<1 M urea, 20 mM EPPS, pH 8.2, 37°C, 16 h). Digests were labeled using 16-plex tandem mass tag (TMT16pro Thermo Fisher Scientific) according to the manufacturer’s instructions, pooled and desalted using C18 columns. Digested and labelled peptide samples were then dried and stored at - 80°C until, depending on the sample complexity, either subjected directly to liquid chromatography tandem mass spectrometry (LC-MS/MS) analysis or first fractionated off-line as described below.

Fractionation by High pH Reversed Phase Chromatography. Peptides were fractionated as described previously (1, 16). Briefly, a Dionex Ultimate 3000 system (Thermo Scientific, Germany) with a Xbridge Peptide BEH C18 column (length, 25cm; inner diameter, 2.1 mm; particle size, 3.5 μm; pore size, 300 Å; Waters) was used. Fractionation was applied via a 50 min gradient solvent system consisting of 20 mM NH4OH in H2O (solvent A) and 20 mM NH4OH in acetonitrile (solvent B), with a flow rate of 200 µL/min. Fractionation was monitored by UV absorbance at 214 nm. A total of 96 fractions of 100 µL each were collected and concatenated into 24 fractions for CLMC with TNFα added or 48 fractions for LPS treated cells. Fractions were dried and stored at - 80°C until LC-MS/MS analysis.

### LC−MS/MS Analysis

Samples were analyzed as previously described (1, 16). Briefly, analyses were performed using an UltiMate 3000 nanoUPLC system connected online to an Exploris 480 or a Lumos Orbitrap mass spectrometer equipped with an EASY nanoSpray source (all - Thermo Fisher Scientific). Peptide separation was performed using an EASY-Spray C18 reversed-phase nano LC column (Acclaim PepMap RSLC; length, 50 cm; inner diameter, 2 μm; particle size, 2 μm; pore size, 100 Å; Thermo Fisher Scientific) at 55°C with a flow rate of 300 nL/min. Peptides were separated using a binary solvent system consisting of 0.1% formic acid, 2% acetonitrile (solvent A) and 0.1% FA and 98% acetonitrile (solvent B). Peptides were eluted by the gradient 2−27% B, followed by a ∼15 min washing step (27−95% B), and a ∼10 min re-equilibration step (2% B). Depending on sample complexity, the total running time varied between 120 min, 150 min and 240 min. Mass spectra were acquired in the positive ion mode with a mass to charge (m/z) range of 375−1500 and a nominal resolution of 120,000 at m/z 200. MS/MS spectra were obtained by data dependent acquisition with dynamic exclusion time of previously selected precursor ions of 45 s. Most abundant peptide ions were selected for higher-energy collision dissociation (HCD) with a normalized collision energy value set at 33 or 35%. The tandem MS (MS/MS) spectra were acquired at a nominal resolution of 45,000 or 50,000. The first mass was fixed at 100 m/z, and the isolation window was 1.6 m/z units.

### Protein Identification and Quantification

The raw LC-MS/MS data were analyzed by Proteome Discoverer (PD, version 3.1) with TMTpro isobaric labeling mass tag. Using Sequest search engine, the MS/MS data were matched against the Uniprot human proteome reference database (Taxonomy ID 9606, 20,306 entries, updated November 2023) supplemented with 1) the INX-heavy and INX-light chains, 2) the sequences of the internal standards trypsinogen (Bo taurus) and myoglobin (Equus caballus) and 3) a common contaminant database. Reversed sequences of the above were concatenated for controlling the false discovery rate (FDR). Peptide mass error tolerance was set at 10 ppm, while MS/MS fragment mass accuracy was set at 0.02 Da. Cysteine carbamidomethylation was used as a fixed modification; TMT-related modifications, methionine oxidation, asparagine and glutamine deamidation were used as variable modifications. Trypsin was selected as enzyme specificity with a maximum of two missed cleavages. Percolator (PD node) with 1% FDR was used as a filter. After removing contaminants, only proteins above the FDR threshold with at least one unique peptide and quantified with at least two peptides were included in the final data set.

### Statistical Analysis

Protein abundances for the background experiment were normalized to the internal standards and adjusted to the difference in added sample volume (i.e. 1/3 (samples: 0, 0.125 and 0.05 mg/mL), 1/2 (samples: 0, 0.125 and 0.05 mg/mL) and 1/1 (samples: 0.8 mg/mL)), respectively. Protein abundances for cell lysates and intact cells were normalized to the total protein abundance in the respective sample. For all samples, the abundance of each protein or peptide was then scaled by dividing it by the average abundance in the control samples. For the background experiment, univariate statistical analysis was performed for normalized protein abundance using two-tailed Student’s t test, with the significance threshold set at 0.05. For the proteome-level experiments, at first peptide abundances were evaluated using Student’s t test. Then p-values of the top 3 most significant peptides for each protein were then combined according to Fisher’s method (18). The FDR values for proteins were corrected for multiple hypotheses according to the Storey Tibshirani method (27). Final protein ranking was done based on the absolute value of the log2 fold-change of the protein abundance between the test groups multiplied by -log10 of the corrected p-values of significant proteins.

## Author contributions

Conceptualization: SLL, RAZ and MG. Methodology: WL, YH, SM, and SLL. Investigation: WL, SLL, RAZ and MG. Visualization: WL and SLL. Resources: RAZ, MG and JZH. Supervision: SLL, SM, RAZ, MG and JZH. Writing—original draft: SLL, WL and RAZ. Writing—review & editing: WL, SLL, YH, SM, JZH, MG and RAZ.

## References

1. Gaetani M, Sabatier P, Saei AA, Beusch CM, Yang Z, Lundstrom SL, et al. Proteome Integral Solubility Alteration: A High-Throughput Proteomics Assay for Target Deconvolution. J Proteome Res. 2019;18(11):4027–37.

2. George AL, Duenas ME, Marin-Rubio JL, Trost M. Stability-based approaches in chemoproteomics. Expert Rev Mol Med. 2024;26:e6.

3. Maity R, Zhang X, Liberati FR, Scribani Rossi C, Cutruzzola F, Rinaldo S, et al. Merging multi-omics with proteome integral solubility alteration unveils antibiotic mode of action. Elife. 2024;13.

4. Knyazeva A, Li S, Corkery DP, Shankar K, Herzog LK, Zhang X, et al. A chemical inhibitor of IST1-CHMP1B interaction impairs endosomal recycling and induces noncanonical LC3 lipidation. Proc Natl Acad Sci U S A. 2024;121(17):e2317680121.

5. Ducellier S, Demeules M, Letribot B, Gaetani M, Michaudel C, Sokol H, et al. Dual molecule targeting HDAC6 leads to intratumoral CD4+ cytotoxic lymphocytes recruitment through MHC-II upregulation on lung cancer cells. J Immunother Cancer. 2024;12(4).

6. Goulet DR, Atkins WM. Considerations for the Design of Antibody-Based Therapeutics. J Pharm Sci. 2020;109(1):74–103.

7. Koch KC, Tew GN. Functional antibody delivery: Advances in cellular manipulation. Adv Drug Deliv Rev. 2023;192:114586.

8. Zhang C, Otjengerdes RM, Roewe J, Mejias R, Marschall ALJ. Applying Antibodies Inside Cells: Principles and Recent Advances in Neurobiology, Virology and Oncology. BioDrugs. 2020;34(4):435–62.

9. Kalliolias GD, Ivashkiv LB. TNF biology, pathogenic mechanisms and emerging therapeutic strategies. Nat Rev Rheumatol. 2016;12(1):49–62.

10. Idriss HT, Naismith JH. TNF alpha and the TNF receptor superfamily: structure-function relationship(s). Microsc Res Tech. 2000;50(3):184–95.

11. van Loo G, Bertrand MJM. Death by TNF: a road to inflammation. Nat Rev Immunol. 2023;23(5):289–303.

12. Holbrook J, Lara-Reyna S, Jarosz-Griffiths H, McDermott M. Tumour necrosis factor signalling in health and disease. F1000Res. 2019;8.

13. Eissner G, Kirchner S, Lindner H, Kolch W, Janosch P, Grell M, et al. Reverse signaling through transmembrane TNF confers resistance to lipopolysaccharide in human monocytes and macrophages. J Immunol. 2000;164(12):6193–8.

14. Kohno T, Tam LT, Stevens SR, Louie JS. Binding characteristics of tumor necrosis factor receptor-Fc fusion proteins vs anti-tumor necrosis factor mAbs. J Investig Dermatol Symp Proc. 2007;12(1):5–8.

15. Ebert EC. Infliximab and the TNF-alpha system. Am J Physiol Gastrointest Liver Physiol. 2009;296(3):G612–20.

16. Zhang X, Lytovchenko O, Lundstrom SL, Zubarev RA, Gaetani M. Proteome Integral Solubility Alteration (PISA) Assay in Mammalian Cells for Deep, High-Confidence, and High-Throughput Target Deconvolution. Bio Protoc. 2022;12(22).

17. Grabert K, Engskog-Vlachos P, Skandik M, Vazquez-Cabrera G, Murgoci AN, Keane L, et al. Proteome integral solubility alteration high-throughput proteomics assay identifies Collectin-12 as a non-apoptotic microglial caspase-3 substrate. Cell Death Dis. 2023;14(3):192.

18. Fisher RA. Statistical Methods for Research Workers, 4th edition 1932.

19. Reichhardt MP, Loimaranta V, Lea SM, Johnson S. Structures of SALSA/DMBT1 SRCR domains reveal the conserved ligand-binding mechanism of the ancient SRCR fold. Life Sci Alliance. 2020;3(4).

20. Holmskov U, Thiel S, Jensenius JC. Collections and ficolins: humoral lectins of the innate immune defense. Annu Rev Immunol. 2003;21:547–78.

21. Good DM, Zubarev RA. Drug target identification from protein dynamics using quantitative pathway analysis. J Proteome Res. 2011;10(5):2679–83.

22. Wang HH, Tsourkas A. Cytosolic delivery of inhibitory antibodies with cationic lipids. Proc Natl Acad Sci U S A. 2019;116(44):22132–9.

23. Pardridge WM. Receptor-mediated drug delivery of bispecific therapeutic antibodies through the blood-brain barrier. Front Drug Deliv. 2023;3.

24. Karimi M, Aslanabadi A, Atkinson B, Hojabri M, Munawwar A, Zareidoodeji R, et al. Subcutaneous liposomal delivery improves monoclonal antibody pharmacokinetics in vivo. Acta Biomater. 2025;195:522–35.

25. Saei AA, Lundin A, Lyu H, Gharibi H, Luo H, Teppo J, et al. Multifaceted Proteome Analysis at Solubility, Redox, and Expression Dimensions for Target Identification. Adv Sci (Weinh). 2024;11(38):e2401502.

26. Feliu N, Walter MV, Montanez MI, Kunzmann A, Hult A, Nystrom A, et al. Stability and biocompatibility of a library of polyester dendrimers in comparison to polyamidoamine dendrimers. Biomaterials. 2012;33(7):1970–81.

27. Storey JD, Tibshirani R. Statistical significance for genomewide studies. Proc Natl Acad Sci U S A. 2003;100(16):9440–5.

