## Supplemental Tables 1 to 3 for "Proteome integral solubility alteration assay for monitoring antibody-antigen cell interactions"

Weiqi Lu *et al.*

**This PDF file includes:**

Tables S1 to S3

Table S1. Peptides from endogenous TNFα in the LPS treaded cells. DG: Deglycosylated, S: soluble, M: membrane

|  |  | |  | **INX** |  | **DG-INX** |  | **Fab INX** |  |  |
| --- | --- | --- | --- | --- | --- | --- | --- | --- | --- | --- |
|  |  | **Peptides** | | **p** | **change** | **p** | **change** | **p** | **change** | **Part of sequence** |
| Lysate |  | [K].VNLLSAIK.[S] | | 1E-06 | 3.8 | 2E-06 | 3.4 | 7.1E-07 | 4.0 | S |
|  |  | [R].LSAEINRPDYLDFAESGQVYFGIIAL.[-] | | 6E-06 | 1.7 | 5E-03 | 1.7 | 8.7E-06 | 1.8 | S |
|  |  | [R].TPSDKPVAHVVANPQAEGQLQWLNR.[R] | | 3E-05 | 2.3 | 2E-04 | 1.8 | 5.8E-04 | 1.9 | S |
|  |  | [R].ANALLANGVELR.[D] | | 4E-05 | 2.4 | 2E-05 | 2.1 | 3.2E-05 | 2.4 | S |
|  |  | [R].ANALLANGVELR.[D] | | 5E-05 | 2.4 | 5E-05 | 2.3 | 1.5E-05 | 2.6 | S |
|  |  | [R].DNQLVVPSEGLYLIYSQVLFK.[G] | | 7E-05 | 1.5 | 3E-02 | 1.5 | 9.3E-04 | 1.6 | S |
|  |  | [R].IAVSYQTK.[V] | | 1E-04 | 1.9 | 7E-04 | 1.7 | 1.9E-04 | 1.8 | S |
|  |  | [R].DNQLVVPSEGLYLIYSQVLFK.[G] | | 2E-04 | 3.5 | 6E-04 | 3.2 | 2.4E-03 | 4.3 | S |
|  |  | [R].TPSDKPVAHVVANPQAEGQLQWLNR.[R] | | 5E-04 | 2.7 | 1E-04 | 1.8 | 1.7E-02 | 2.6 | S |
|  |  | [R].TPSDKPVAHVVANPQAEGQLQWLNR.[R] | | 6E-04 | 1.4 | 1E-02 | 1.2 | 1.6E-03 | 1.4 | S |
|  |  | [R].TPSDKPVAHVVANPQAEGQLQWLNRR.[A] | | 2E-03 | 2.0 | 1E-02 | 1.6 | 9.2E-03 | 1.6 | S |
|  |  | [K].GQGCPSTHVLLTHTISR.[I] | | 2E-02 | 1.9 | 7E-03 | 1.5 | 1.2E-03 | 1.7 | S |
|  |  | [R].DNQLVVPSEGLYLIYSQVLFKGQGCPSTHVLLTHTISR.[I] | | 6E-02 | 1.5 | 5E-01 | 1.1 | 5.6E-01 | 1.1 | S |
|  |  | [R].DVELAEEALPK.[K] | | 9E-02 | 1.3 | 1E-01 | 1.2 | 1.5E-01 | 1.2 | M |
|  |  | [R].IAVSYQTK.[V] | | 3E-01 | 0.8 | 3E-01 | 0.8 | 4.3E-01 | 0.8 | S |
|  |  | [K].VNLLSAIK.[S] | | 3E-01 | 1.2 | 7E-01 | 1.1 | 2.4E-01 | 1.3 | S |
|  |  | [R].TPSDKPVAHVVANPQAEGQLQWLNRR.[A] | | 4E-01 | 1.4 | 9E-01 | 1.1 | 7.7E-01 | 1.1 | S |
|  |  | [R].DNQLVVPSEGLYLIYSQVLFK.[G] | | 5E-01 | 0.6 | 6E-01 | 0.6 | 6.0E-01 | 0.6 | S |
|  |  | [R].TPSDKPVAHVVANPQAEGQLQWLNR.[R] | | 6E-01 | 0.8 | 4E-01 | 0.7 | 6.5E-01 | 0.8 | S |
| Intact cell | | [R].ANALLANGVELR.[D] | | 1E-06 | 0.4 | - | - | 2E-06 | 0.4 | S |
|  |  | [K].VNLLSAIK.[S] | | 2E-06 | 0.4 | - | - | 2E-06 | 0.4 | S |
|  |  | [R].ANALLANGVELR.[D] | | 6E-04 | 0.7 | - | - | 2E-03 | 0.8 | S |
|  |  | [R].DVELAEEALPK.[K] | | 4E-02 | 0.9 | - | - | 2E-02 | 0.9 | M |

Table S2. Top 0.5% of proteins from the PISA-AAI experiment in lysed LPS-treated cells using INX. Only two proteins, TEP1 and TNF, were also among the top 0.5% when the same experiment was performed with the Fab fragment.

| Top 0.5% | INX+LMCL |  | | | | Fab-INX+LMCL | | | |
| --- | --- | --- | --- | --- | --- | --- | --- | --- | --- |
| **Protein** | **log 10(p)** | **log2 (INX/C)** | **score** | **rank** | **log 10(p)** | | **log2 (INX/C)** | **score** | **rank** |
| CPLANE2 | 3.7 | 4.22 | 15.8 | 1 | 0.2 | | 0.1 | - | - |
| PTGIR | 3.0 | 3.47 | 10.5 | 2 | 0.6 | | 0.4 | - | - |
| APOA2 | 5.2 | 1.82 | 9.4 | 3 | 0.7 | | 1.6 | - | - |
| NBEAL1 | 4.0 | 2.23 | 9.0 | 4 | 0.3 | | 0.2 | - | - |
| ALB | 9.0 | 0.99 | 8.9 | 5 | 3.7 | | 0.7 | 3.5 | 124 |
| TF | 6.5 | 1.35 | 8.8 | 6 | 2.2 | | 1.2 | 2.6 | 684 |
| APOA1 | 6.0 | 1.41 | 8.5 | 7 | 1.9 | | 1.2 | 2.2 | 1120 |
| **TNF** | **7.4** | **0.80** | **5.9** | **8** | **4.8** | | **0.80** | **5.4** | **2** |
| IGHA1 | 5.3 | 0.93 | 4.9 | 9 | 2.2 | | 0.7 | 2.4 | 885 |
| A2M | 5.5 | 0.70 | 3.9 | 10 | 2.3 | | 0.6 | 2.5 | 808 |
| GC | 5.4 | 0.69 | 3.7 | 11 | 2.3 | | 0.4 | 2.3 | 1101 |
| FAM104A | 2.2 | 1.69 | 3.7 | 12 | 2.3 | | 1.5 | 2.6 | 722 |
| FBXO46 | 2.9 | 1.25 | 3.7 | 13 | 2.5 | | 1.1 | 2.8 | 448 |
| SDF2 | 4.1 | 0.86 | 3.5 | 14 | 1.5 | | 0.3 | 1.4 | 2807 |
| EML6 | 1.9 | 1.65 | 3.2 | 15 | 2.2 | | 1.8 | 2.9 | 418 |
| UQCC2 | 3.8 | 0.80 | 3.1 | 16 | 1.3 | | 0.0 | 1.0 | 3143 |
| NDUFS5 | 2.6 | -1.13 | 2.9 | 17 | 1.0 | | -1.2 | - | - |
| PRPSAP1 | 4.2 | 0.69 | 2.9 | 18 | 0.9 | | 0.2 | - | - |
| EIF1AX | 3.1 | -0.91 | 2.8 | 19 | 3.9 | | -1.0 | 2.9 | 397 |
| SLC16A3 | 3.0 | 0.94 | 2.8 | 20 | 1.8 | | 0.7 | 1.8 | 1834 |
| GNB1L | 3.7 | 0.72 | 2.7 | 21 | 1.5 | | 0.6 | 1.6 | 2360 |
| USP48 | 5.3 | 0.49 | 2.6 | 22 | 0.3 | | 0.2 | - | - |
| IGHM | 3.6 | 0.70 | 2.5 | 23 | 1.0 | | 0.7 | - | - |
| SYTL3 | 3.1 | 0.80 | 2.5 | 24 | 1.9 | | 1.2 | 2.5 | 764 |
| PPP1R15B | 3.5 | 0.71 | 2.5 | 25 | 0.3 | | 0.0 | - | - |
| FCRL3 | 2.1 | 1.11 | 2.4 | 26 | 2.0 | | 1.0 | 2.1 | 1370 |
| STX5 | 3.9 | 0.60 | 2.3 | 27 | 1.7 | | 0.2 | 1.5 | 2550 |
| TPRA1 | 1.9 | 1.17 | 2.3 | 28 | 2.1 | | 1.3 | 2.6 | 632 |
| CTNNBIP1 | 2.6 | 0.83 | 2.2 | 29 | 1.3 | | 0.7 | - | - |
| DAPK1 | 3.0 | 0.69 | 2.1 | 30 | 2.1 | | 0.6 | 2.2 | 1194 |
| **TEP1** | **5.5** | **0.39** | **2.1** | **31** | **4.8** | | **0.2** | **4.6** | **5** |
| FLII | 5.6 | 0.37 | 2.1 | 32 | 2.9 | | 0.3 | 2.9 | 435 |
| ORM1 | 2.1 | 0.98 | 2.1 | 33 | 1.3 | | 0.9 | 1.5 | 2562 |
| BANF1 | 1.7 | -1.22 | 2.1 | 34 | 2.9 | | -1.6 | 1.7 | 2079 |
| MIER3 | 2.4 | 0.85 | 2.0 | 35 | 0.7 | | 0.4 | - | - |
| KDM1A | 4.8 | 0.42 | 2.0 | 36 | 2.7 | | 0.3 | 2.6 | 646 |
| ANKIB1 | 2.5 | 0.80 | 2.0 | 37 | 1.8 | | 0.6 | 1.9 | 1682 |
| GDAP1 | 2.7 | 0.72 | 2.0 | 38 | 2.6 | | 0.9 | 3.1 | 305 |
| BASP1 | 2.03 | -0.97 | 1.96 | 39 | 2.89 | | -0.97 | 2.11 | 1339 |

Table S3. Top 0.5% of proteins from the PISA-AAI experiment on intact LPS-treated cells that overlapped between experiments with the intact antibody and the Fab fragment.

| **Top 0.5% overlap** | INX+LMCL | | | | Fab-INX+LMCL | | | |  | Ranked according to: |
| --- | --- | --- | --- | --- | --- | --- | --- | --- | --- | --- |
| **Protein** | **log 10(p)** | **INX/C** | **score** | **rank** | **log 10(p)** | **log2 (INX/C)** | **score** | **rank** | | **log2[INX/C]/log2[Fab-INX/C]** |
| DMBT1 | 6 | -1.6 | 9.68 | **4** | 5.6 | -1.1 | 6.33 | 16 | | 1.38 |
| TNF | 10 | -0.9 | 8.53 | **8** | 9.0 | -0.8 | 7.09 | 13 | | 1.20 |
| STX10 | 7 | -1.9 | 13.09 | **2** | 7.0 | -1.8 | 12.81 | 3 | | 1.04 |
| F10 | 5 | -1.6 | 7.67 | **10** | 4.5 | -1.5 | 6.83 | 15 | | 1.02 |
| IGFBP2 | 7 | -1.0 | 6.66 | **13** | 5.8 | -1.0 | 5.83 | 19 | | 1.01 |
| GP1BA | 9 | -0.8 | 6.78 | **12** | 7.8 | -0.8 | 5.99 | 17 | | 0.99 |
| ITGA2B | 11 | -0.4 | 4.92 | **39** | 10.7 | -0.5 | 4.90 | 34 | | 0.97 |
| MPIG6B | 7 | -0.8 | 5.40 | **27** | 6.5 | -0.8 | 5.44 | 25 | | 0.96 |
| IGF2 | 4 | -1.5 | 5.40 | **26** | 3.6 | -1.6 | 5.62 | 21 | | 0.93 |
| RBP4 | 10 | -1.3 | 13.30 | **1** | 9.0 | -1.4 | 12.96 | 2 | | 0.93 |
| MAPT | 9 | 0.6 | 5.05 | **35** | 7.4 | 0.6 | 4.76 | 38 | | 0.92 |
| COL6A1 | 9 | -1.1 | 9.22 | **6** | 8.7 | -1.2 | 10.50 | 6 | | 0.89 |
| CD163 | 12 | -0.7 | 9.23 | **5** | 12.0 | -0.8 | 10.15 | 7 | | 0.88 |
| COMP | 9 | -1.0 | 8.77 | **7** | 8.8 | -1.1 | 9.85 | 8 | | 0.87 |
| PTPN1 | 13 | -0.4 | 5.28 | **30** | 11.8 | -0.5 | 5.56 | 23 | | 0.87 |
| AFP | 4 | -1.4 | 6.22 | **15** | 4.4 | -1.7 | 7.45 | 10 | | 0.85 |
| IGFBP5 | 5 | -0.9 | 4.89 | **41** | 5.4 | -1.1 | 5.98 | 18 | | 0.83 |
| ANXA5 | 12 | -1.0 | 12.92 | **3** | 12.0 | -1.3 | 15.80 | 1 | | 0.79 |
| AOC3 | 4 | -1.3 | 5.52 | **21** | 4.3 | -1.7 | 7.30 | 12 | | 0.76 |
| ELANE | 10 | -0.6 | 5.75 | **18** | 9.0 | -0.8 | 7.02 | 14 | | 0.75 |
| ANXA1 | 11 | -0.7 | 7.44 | **11** | 11.1 | -1.0 | 11.28 | 5 | | 0.68 |
| ANXA4 | 12 | -0.7 | 7.90 | **9** | 12.3 | -1.0 | 12.28 | 4 | | 0.65 |
